# Synthetic Echocardiograms Generation Using Diffusion Models

**DOI:** 10.1101/2023.11.11.566718

**Authors:** Alexandre Olive Pellicer, Amit Kumar Singh Yadav, Kratika Bhagtani, Ziyue Xiang, Zygmunt Pizlo, Irmina Gradus-Pizlo, Edward J. Delp

**Affiliations:** Video and Image Processing Lab (VIPER), School of Electrical and Computer Engineering, Purdue University, West Lafayette, Indiana, USA

**Keywords:** chocardiograms, diffusion model, generative AI

## Abstract

An echocardiogram is a video sequence of a human heart captured using ultrasound imaging. It shows heart structure and motion and helps in diagnosis of cardiovascular diseases. Deep learning methods, which require large amounts of training data have shown success in using echocardiograms to detect cardiovascular disorders such as valvular heart disease. Large datasets of echocardiograms that can be used for machine learning training are scarce. One way to address this problem is to use modern machine learning generative methods to generate synthetic echocardiograms that can be used for machine learning training. In this paper, we propose a video diffusion method for the generation of echocardiograms. Our method uses a 3D selfattention mechanism and a super-resolution model. We demonstrate that our proposed method generates echocardiograms with higher resolution and with lesser artifacts, compared to existing echocardiogram generation methods.

## I. Introduction

Deep learning methods have been successful in many biomedical applications such as disease diagnosis [1]–[4], protein structure prediction [5], and image reconstruction from brain activity [6]. Several deep learning methods [7]–[9] have been used to examine echocardiograms to assist clinicians in identifying cardiac abnormalities and diagnosing cardiovascular diseases. Echocardiograms are ultrasound sequences that capture the structure and motion of the heart [10] (see Figure 1).

**Fig. 1.**
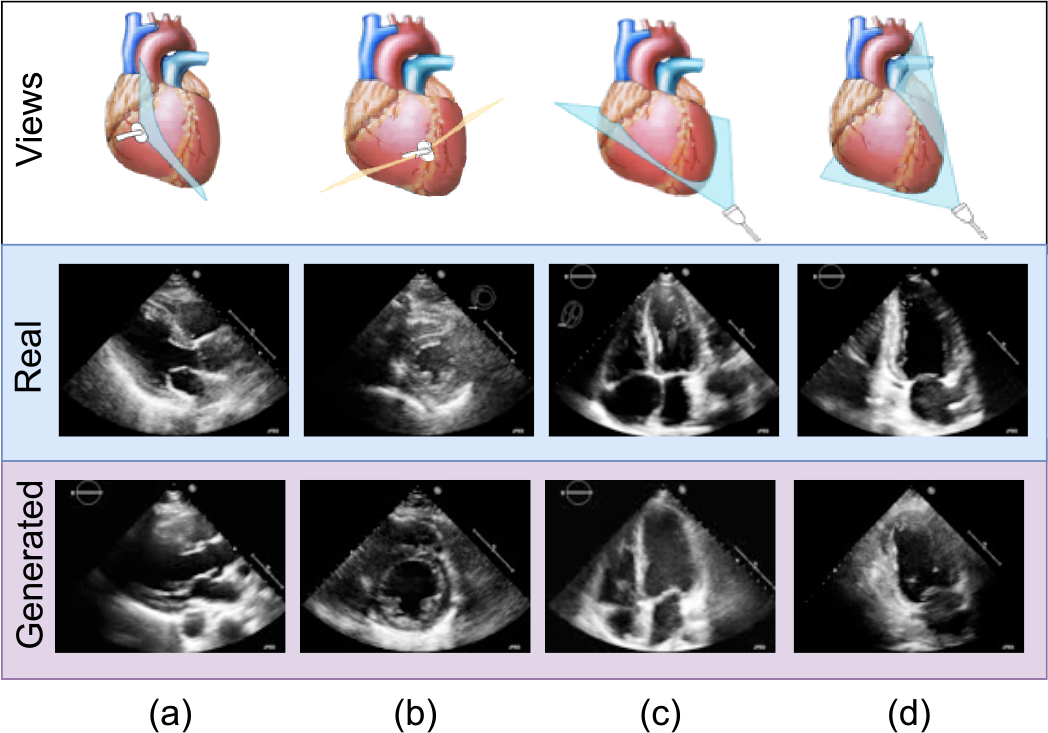
Echocardiograms from 4 different views of the heart. (a) Parasternal long axis view; (b) Parasternal short axis view; (c) Apical 4-chamber view; (d) Apical 2-chamber view. The position of the ultrasound transducer is shown in the top row. The middle row shows real echocardiograms corresponding to the transducer position. The bottom row shows the echocardiograms generated by our method.

Self-supervised deep learning methods learn feature representations from unlabeled data by using pretext tasks such as reconstruction of input data from the learned feature representations [11]. Such methods have shown promising results in detecting abnormalities [12] in medical images such as eye diseases [13], and cardiac ejection fraction prediction [14]. These methods need large amounts of unlabeled data (at a scale of thousands or even millions of images) [15].

For machine learning methods to achieve optimal classification performance a large number of examples with known pathology needs to be available. The required scale is in thousands or even millions of examples. In the world of echocardiography those examples exist but their use is limited by the need to de-identify patient specific information due to patient privacy regulations and by the fact that datasets are fragmented by storage in individual hospital or clinic databases.

Once echocardiogram deep learning methods are developed their use will be revolutionary. From teaching technicians and physicians to obtain proper images in simulated environment to algorithm based acquisition of optimal images in patient care to performing automated measurements of echo parameters which currently doubles the time the technician needs to perform a study to automated interpretation of studies. In practice, this makes the real datasets not accessible for training self-supervised methods.

One way to address the problems above is to use synthetic data for echocardiogram analysis. Generative machine learning methods, such as GANs [16] or diffusion models [17] have been used to generate synthetic data that can be used for training [18] 22]. In this paper, we propose a video diffusion method to generate echocardiograms. We also experiment with Real-Enhanced Super-Resolution Generative Adversarial Network (Real-ESRGAN) [23], a super-resolution method which increases the resolution of each frame of the echocardiogram.

The main contributions of this work are: a) we propose a modified video diffusion model that can generate echocardiograms; b) we propose a self-attention approach and investigate different noise schedules to improve the performance of our method; c) we investigate Real-ESRGAN as a super-resolution approach.

It should be noted that we are not generating echocardiograms with ground truth annotations (e.g., heart wall boundaries). The sequences we generate can be used for selfsupervised learning approaches we discussed above.

## II. Related Work

Diffusion models have been used to generate images [18], [24], [25], and speech signals [26]. Recently, several methods have been proposed which explore the generation of video sequences [27]–[29]. Hong *et al*. proposed CogVideo [27], a text-to-video generation transformer which fine-tuned a pretrained text-to-image generation method CogView-2 [28]. Ho *et al*. proposed a Video Diffusion Model (VDM) trained using image and video data which was used for the synthesis of videos with a resolution of 64 × 64 pixels [30]. These low-resolution videos were up sampled to 128 × 128 pixels using super-resolution methods [30]. Super-resolution methods enhance the resolution of an image generally using convolutional up sampling layers [23]. Singer *et al*. proposed MakeA-Video, a diffusion model trained on text-image pairs and unsupervised videos for video generation [29]. The authors also proposed super-resolution approaches to enhance the resolution of generated videos [29].

Diffusion models have been proposed for generating echocardiograms [18], [31]. Stojanovski *et al*. [18] proposed “Echo from Noise” to produce one annotated image of an echocardiogram. This method does not produce echocardiogram sequences. The annotations are segmentation masks indicating the left ventricle and left atrium of the heart. Several of these generated still images were used as training data for networks to segment regions of the heart from frames of echocardiogram sequences. Echo from Noise uses a semantic diffusion model [32] to generate synthetic images conditioned on semantic labels of the heart’s structure.

Methods for generating echocardiogram sequences have also been proposed. Reynaud *et al*. used video diffusion models to generate synthetic sequences of echocardiograms, which can be used for training machine learning methods and instructing medical professionals in the interpretation of echocardiograms [31]. The authors used a cascade diffusion model [33] for video-prediction. In this method, an initial frame of an actual echocardiogram is required, along with the Left Ventricular Ejection Fraction (LVEF)1, to generate the rest of the echocardiogram sequence. The generated echocardiogram sequences are not annotated with respect to structures such as heart walls but each generated echocardiogram has the LVEF associated with it. Echocardiograms can be captured from different views of the heart as shown in Figure 1. The method proposed by Reynaud *et al.* [31] is limited to the generation of echocardiograms from the Apical 4-chamber view (see Figure 1). Our proposed method can generate higher resolution echocardiograms and from four different views which are shown in Figure 1.

## III. Proposed Method

The block diagram of our proposed method is shown in Figure 2 and consists of two parts. First, a video diffusion model [30] is used to generate echocardiogram sequences. Then, a super-resolution method [23] is used to increase the spatial resolution of the frames in the sequence. This section contains a detailed description of each part.

**Fig. 2.**
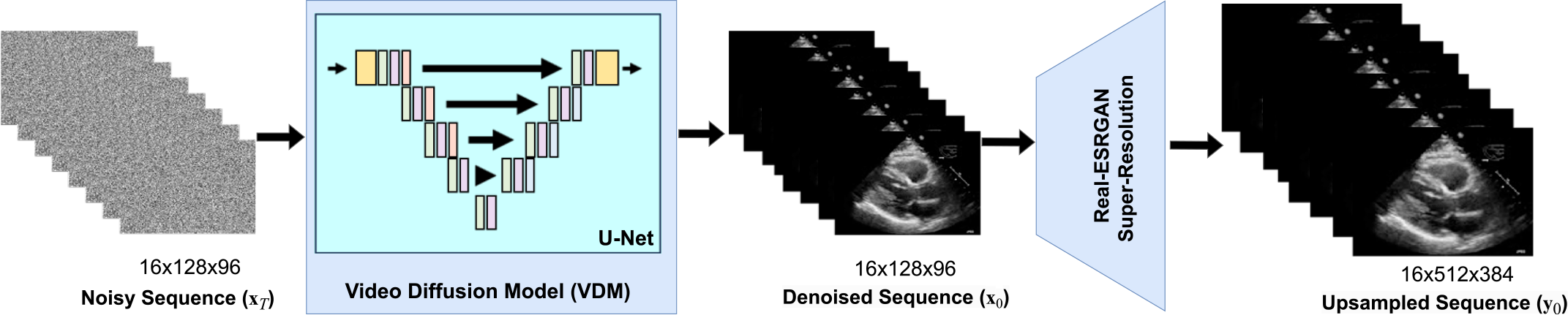
Block diagram of our proposed method.

### A. Initial Sequence Generation

We use a denoising probabilistic diffusion model [34]–[36] which uses a parameterized Markov chain structure to characterize the distribution of the pixels in the sequence *q* (*x*_0_), where x_0_ denotes pixels in an echocardiogram sequence. Given a distribution x_0_ ∼ *q* (x_0_), we define a forward noising process *q* which produces x_*t*_, (*t* ∈ {1, 2…, *T*}) by adding Gaussian noise at diffusion step *t* (*T* denotes the total number of diffusion steps) with variance 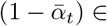 (0, 1) as follows:

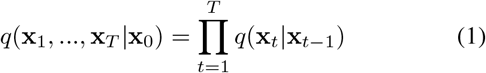

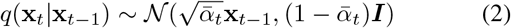

Given sufficiently large *T* and a well behaved noise schedule of 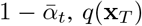 is nearly an isotropic Gaussian distribution. Therefore, if the reverse distribution *q*(*x*_*t*−1_ | *x*_*t*_) is known, a x_0_ can be sampled from the distribution *q* (*x*_0_) by starting with *x*_*T*_ ∼ 𝒩 (0, **I**) and iteratively denoising *x*_*T*_. The Gaussian noise x_T_ is iteratively denoised to obtain less noisy sequences *x*_*T* −1_, *x*_*T* −2_, …, *x*_0_. The sequence obtained after the last denoising step i.e., *x*_0_ is the generated echocardiogram. The reverse distribution *q*(*x*_*t*−1_ | *x*_*t*_) depends on the distribution of training data, in our case the distribution of echocardiogram sequences. Therefore, *q*(*x*_*t*−1_|*x*_*t*_) is estimated using a network as *p*_*θ*_(*x*_*t*−1_ | *x*_*t*_). *p*_*θ*_(*x*_*t*−1_ | *x*_*t*_) and *θ* denote the distribution estimated by the network and parameters of the network, respectively.

In previous work on image generation using diffusion models [35], [36], the network used for estimating *p*_θ_(x_*t*−1_ | x_*t*_) consists of a U-Net [37] as shown in Figures 2 and 3. The U-Net is a type of Convolutional Neural Network (CNN) that consists of several encoder-decoder pairs. Each pair is known as a layer (see Figure 3), where the encoder down samples, and the decoder up samples. There are skip connections [38] to connect the encoder and decoder blocks as shown in Figure 3. To generate sequences of echocardiograms, we use the modified U-Net for spatiotemporal sequences as described in [30]. The input of the U-Net is five-dimensional with each dimension representing batch size of the network, number of channels in each frame, number of frames in the sequence and the frame size.

**Fig. 3.**
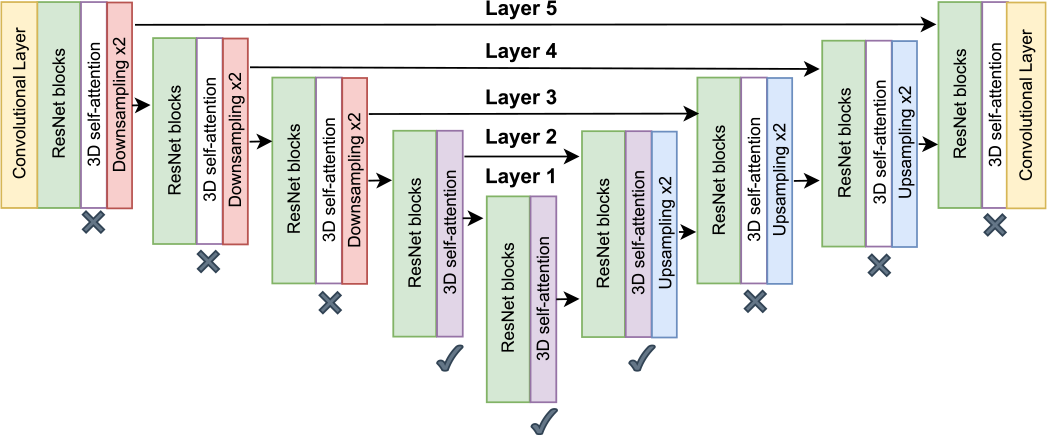
Block diagram of the 3D U-Net and 3D self-attention used in our video diffusion model. The skip-connections are before the down sampling blocks. The ✓ indicates the layers where the self-attention block resulted in the best performance metrics.

Our method incorporates a 3D (two spatial dimensions in each frame and time) self-attention mechanism. We do this by adding 3D self-attention blocks in between some of the layers of the U-Net. This is different than in [30] where the 3D self-attention blocks are inserted between every layer. Our approach helps capture the motion of the heart. This is discussed in detail in Section IV. More details about the 3D self-attention block are described in [30].

### B. Super-Resolution

The output of the Video Diffusion Model (VDM) is denoted by x_0_, as shown in Figure 2. x_0_ is the input to the superresolution model which generates frames of higher spatial resolution denoted by y_0_. For super-resolution, we slightly modified Real-ESRGAN [23] to work with rectangular frames.

We fine-tune the modified Real-ESRGAN for echocardiograms. The synthetic echocardiograms generated by the video diffusion model (x_0_) are of resolution 128 × 96 pixels. These frames have their spatial resolution increased by four times, to generate echocardiograms of resolution i.e., 512 × 384 pixels (y_0_).

## IV. Experiments AND Results

### A. Dataset

We use the IPRS dataset, which was provided by the Indiana University School of Medicine. It contains echocardiograms from 50 different patients in the 4 different views (see Figure 1). In total, there are 200 echocardiogram sequences. The spatial resolution varies from 800 × 600 pixels to 640 × 480 pixels, and the aspect ratio of the frames is 4:3 for all sequences. Before training the generator, we pre-process the echocardiograms. All frames are resized to 640 × 480 pixels using down sampling. The sequences are then cropped to 512 × 384 pixels. Then these cropped sequences are down sampled to 128 × 96 pixels. We denote 512 × 384 sequences as the “high resolution” sequences and the 128 × 96 sequences as the “low resolution” sequences.

Since the number of frames vary in the sequences, we only examine the first 16 frames. If an echocardiogram has 32 frames or more, we split it into multiple sequences of 16 frames each. After the pre-processing, we construct two datasets:

- A dataset consisting of 224 echocardiograms of the high resolution sequences. This dataset is used as the ground truth for training the super-resolution method.
- A dataset consisting of 224 echocardiograms of low resolution sequences. This dataset is used to train the video diffusion model and as an input to the superresolution method.

### B. Experimental Settings

The video diffusion model is trained for 125,000 iterations. The optimal number of iterations was experimentally determined. We used a batch size of 4, 1000 diffusion steps equals, and the Adam optimizer [39] with a learning rate of 10^−4^. We used an Exponential Moving Average (EMA) [31], [40] with a decay of 0.995 for more stable training and better performance metrics.

For training the super-resolution model, we first trained the Super-Resolution Network (SRNet) (the generative network component of Real-ESRGAN) for 200,000 iterations for faster convergence. Next, we trained Real-ESRGAN (both the generative network component and the adversarial network component) for 300,000 iterations. The rest of the hyperparameters are the same as described in [23].

### C. Results

We are interested in determining some type of qualitative metric for assessing how well our generated sequences correspond to echocardiograms. One metric we use is the Fréchet Inception Distance (FID) [41]. This will allow us to compare distribution of features of the generated echocardiograms and the real echocardiograms in our dataset of real echocardiograms (*IPRS*). To estimate FID, both real and generated frames are analyzed by a pretrained Inception-v3 network [42] that produces a set of features. The FID score denotes the similarity between the features obtained from the generated and real images.

We also use the Fréchet Video Distance (FVD) [43], which is an extension of FID that generalizes to videos and measures similarity between video features obtained from real and generated videos. The lower the FID for a generated image and the lower the FVD for a generated video sequence, the higher is the visual quality of the generated image and video sequence, respectively.

Experiment 1: In this experiment, we determine which layers of the U-Net should contain 3D self-attention blocks to obtain best performance. We experimented by adding the 3D self-attention block to some of the layers of the U-Net, adding one layer at a time. We started by adding the 3D self-attention block to Layer 1 (see Figure 3), then we added to Layers 1 and 2 and continued adding to the subsequent layers. The results of this experiment are shown in Table I. From Table I, we can observe that adding the 3D self-attention block to Layers 1 and 2 corresponds to the best FID and FVD scores. Therefore, we use this for our next experiments.

**TABLE I.**
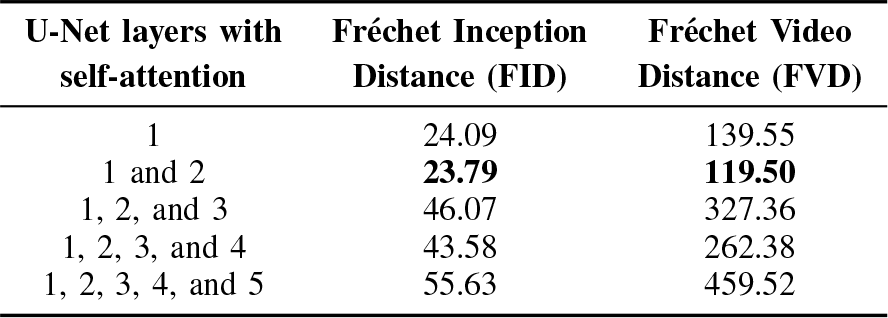
Results of Experiment 1 showing fid and FVD after adding the self-attention block.

Experiment 2: In this experiment, we evaluated the performance of the video diffusion model by varying the noise schedule. We used the self-attention in layer 1 and layer 2 (as shown in Figure 3), which resulted to the best performance in Experiment 1. We used a cosine noise schedule defined as:

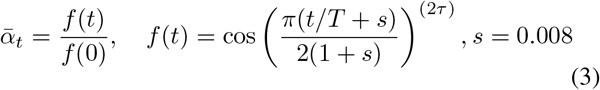

*t* and *T* represent the *t*^th^ diffusion step and the total number of diffusion steps, respectively. Following the experiments described in [44] we analyzed the results obtained when τ is 1, 2, and 3. The results of this experiment are shown in Table II. From Table II, we observe that τ = 2 corresponds to the lowest FID and FVD scores.

**TABLE II.** Results of Experiment 2 showing FID and FVD for different values of τ from Equation 3.

| $\tau$ | Fréchet Inception Distance (FID) | Fréchet Video Distance (FVD) |
| --- | --- | --- |
| 1 | 23.79 | 119.50 |
| 2 | **17.65** | **77.47** |
| 3 | 20.95 | 99.73 |

Experiment 3: The goal of this experiment is to analyze the performance of the super-resolution model. Using the high resolution echocardiogram sequences generated by Real-ESRGAN we computed the FID score of 36.47 and a FVD score of 176.12, respectively.

### D. Comparison with Existing Methods

We first compare our method with the Echo from Noise method proposed by Stojanovski *et al.* [18]. This method is limited to the generation of one 2D echocardiogram image, not a sequence, whereas we generate echocardiogram sequences with cardiac motion [18]. We only use the FID score for comparison here because Echo from Noise only creates a single frame. We used the real and synthetic images provided by [18] to estimate the FID score of the Echo from Noise method. The FID score of echocardiogram images generated using Echo from Noise is 75.36 while FID score of images generated using our method is 17.65. This indicates echocardiograms generated from our method have better quality. A visual comparison of the echocardiograms generated using the Echo from Noise and those using our method is shown in Figure 4. It shows that several details present in the echocardiograms generated from our method are missing in the echocardiograms generated using Echo from Noise.

**Fig. 4.**
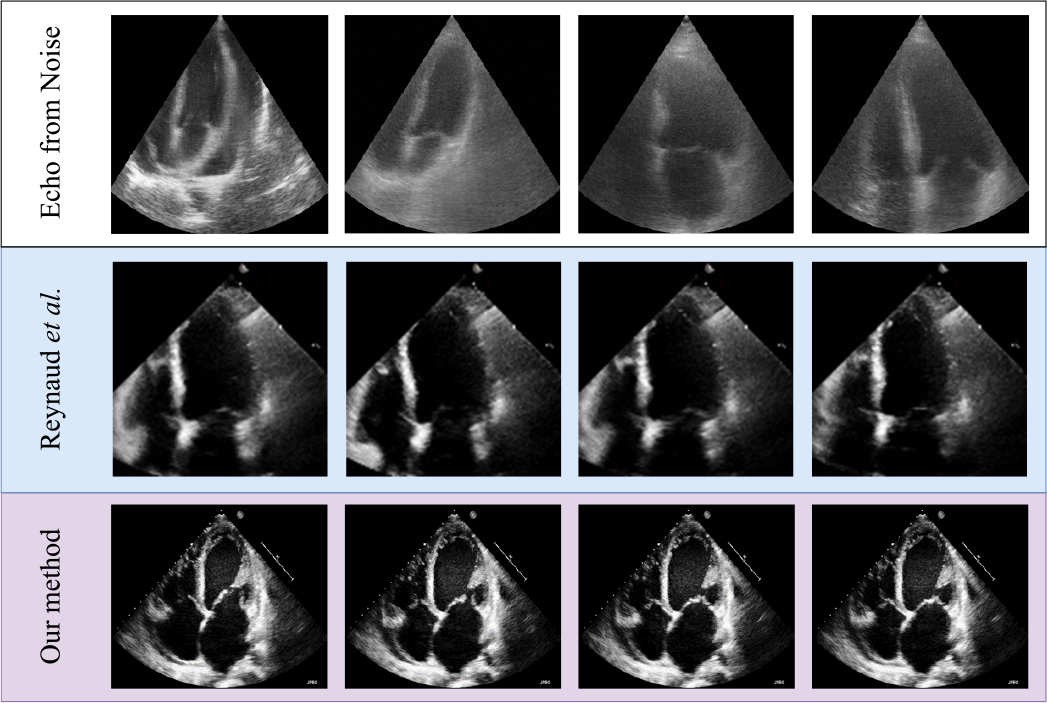
The top row shows echocardiogram frames generated using the Echo from Noise method proposed by Stojanovski *et al.* [18].The middle row shows echocardiogram frames generated using the approach proposed by Reynaud *et al.* [31]. The bottom row shows echocardiogram frames generated using our method.

We also compare our generated echocardiograms with the ones obtained by Reynaud *et al.* [31]. In [31], the authors focused on using video diffusion models for the conditioned generation of lower resolution echocardiograms (112 × 112 pixels) given the first frame and LVEF as input. The *EchoNet-Dynamic* dataset is used [45]. Table III shows a comparison of the *EchoNet-Dynamic* dataset [45] and the *IPRS* dataset we used for our experiments.

**TABLE III.** Comparison between the *EchoNet-Dynamic* dataset [45] and the *IPRS* dataset.

| Dataset | *EchoNet-Dynamic* | *IPRS* |
| --- | --- | --- |
| Number of echocardiograms | 10000 | 224 |
| Format | AVI | AVI |
| Echocardiogram views | Apical 4-chamber | Parasternal long axis<br>Parasternal short axis<br>Apical 4-chamber<br>Apical 2-chamber |
| Frame-resolution | $112 \times 112$ pixels | $512 \times 384$ pixels |

The synthetic echocardiograms generated by Reynaud *et al.* [31] have an FID score of 12.3 and an FVD score of 60.5. These results are obtained for echocardiograms of resolution 112 × 112 pixels. For fair comparison, here we compare our lower resolution echocardiograms (128 × 96 pixels) generated using our video diffusion model with the ones obtained in [31]. Both methods have 16 frames per echocardiogram. The lower resolution echocardiograms generated by our method have an FID score of 17.65 and an FVD score of 77.47 [31]. Note that our method is trained using 44 times less training data.

Although the quantitative metrics for [31] are slightly better than that for our method, our approach can generate four different views. The ultrasound artifacts in our generated sequences appear to be more like artifacts in real echocardiograms. FID, used to assess generated images, is designed for natural images (*i.e.*, people, cars, animals). In these natural images, the goal is to minimize noise. However, in ultrasound images, speckle noise is significant. In medical images, such as echocardiogram images, speckle noise may convey crucial information about pathologies in tissues or organs [46]. For example, methods have been created to identify fibrosis by analyzing speckle noises present in echocardiogram images [47]. Figure 4 shows a comparison of our method, the Echo from Noise method [18], and the method proposed by Reynaud *et al.* [31]. Finally, our approach generates echocardiograms sequences of resolution 512 × 384 pixels, which is higher as compared to both methods, Echo from Noise and the method proposed by Reynaud *et al.*.

Examples of synthetic echocardiograms generated by our method are available : https://alexolpe.github.io/final-thesis/index.html

## V. Conclusion AND Future Work

We proposed a method to generate synthetic echocardiograms of four different views of the heart. We used a video diffusion model and super-resolution to generate echocardiograms. We demonstrated the performance of our method and its capability to generate echocardiograms with more details as compared to existing methods. Future work will focus on generation of annotated echocardiograms. Conditioning the generation of the echocardiograms to characteristics such as types of cardiac abnormalities, wall motion, and ejection fraction will be a focus for future work.

## Footnotes

† The work described in this paper was conducted while the author Olive was affiliated with Escola Tècnica Superior d’Enginyeria de Telecomunicació de Barcelona, Universitat Politècnica de Catalunya (UPC), Barcelona, Spain.

1 LVEF represents the percentage of oxygen-rich blood ejected from the left ventricle during each heart contraction.

## Notes

### Competing Interest Statement

The authors have declared no competing interest.

